# Active transcription regulates ORC/MCM distribution whereas replication timing correlates with ORC density in human cells

**DOI:** 10.1101/778423

**Authors:** Nina Kirstein, Alexander Buschle, Xia Wu, Stefan Krebs, Helmut Blum, Wolfgang Hammerschmidt, Olivier Hyrien, Benjamin Audit, Aloys Schepers

## Abstract

Eukaryotic replication initiates during S phase from origins that have been licensed in the preceding G1 phase. Here, we compare ChIP-seq profiles of the licensing factors Orc2, Orc3, Mcm3, and Mcm7 with replication initiation events obtained by Okazaki fragment sequencing. We demonstrate that MCM is displaced from early replicating, actively transcribed gene bodies, while ORC is mainly enriched at active TSS. Late replicating, H4K20me3 containing initiation zones display enhanced ORC and MCM levels. Furthermore, we find early RTDs being primarily enriched in ORC, compared to MCM, indicating that ORC levels are involved in organizing the temporal order of DNA replication. The organizational connection between active transcription and replication competence directly links changes in the transcriptional program to flexible replication patterns, which ensures the cell’s flexibility to respond to environmental cues.

---

Mammalian DNA replication is a highly orchestrated process ensuring the exact inheritance of genomes of tens to thousands of million base pairs in size. In human cells, replication initiates from 30,000 – 50,000 replication origins per cell^1, 2^. Origins are not activated synchronously but are organized into individual replication timing domains (RTDs), which replicate in a timely coordinated and reproducible order from early to late in S phase^3, 4^. The replication cascade or domino model proposes that within one RTD, replication first initiates at the most efficient origins and then spreads to less efficient origins. RTDs are separated by timing transition regions and it is debated whether replication spreading is blocked at these regions^5, 6^.

The establishment of replication competence occurs in late mitosis and during the G1 phase of the cell cycle^7^. The first step is the cell-cycle dependent assembly of the evolutionary conserved origin recognition complex (ORC) to origins^8, 9^. ORC and two chaperones, Cdt1 and Cdc6, cooperatively load minichromosome maintenance (MCM) complexes as double hexamers^10–12^. The MCM complex is the central unit of the replicative helicase. The resulting multi-subunit complex is termed pre-replicative complex (pre-RC). A single ORC loads multiple MCM helicases, which are translocated from their original loading site, but no ORC, neither Cdc6, nor Cdt1 are required for origin activation^13, 14^.

Replication origins are defined functionally. In the unicellular *S. cerevisiae*, replication origins are genetically characterized by the ARS consensus sequence^15^. In multicellular organisms, replication initiates from flexible locations and no common consensus element for origin selection and activation has yet been identified. It is generally believed that chromatin features including histone modifications, nucleosome dynamics and DNA modifications contribute to origin specification^1, 16, 17^, including H4K20me3 that supports the licensing of a subset of late replicating origins in heterochromatin^18^. Thus, changing environmental conditions, DNA damage and the development status of each cell are integrated into highly dynamic local chromatin profiles, which influence the plasticity of origin selection^1^.

Different approaches have been developed to characterize mammalian replication initiation by single-molecule visualization (DNA combing) or sequencing of purified initiation products (short nascent strands (SNS-seq), initiation site sequencing (INI-seq) replication-bubbles containing restriction fragments (bubble-seq) or elongation intermediates (Okazaki fragments). In various human cell lines, SNS-seq and INI-seq have identified specific replication initiation sites, which mainly correlate with transcriptional start sites (TSS) and locate close to CG-rich regions that are enriched with G-quadruplex motifs (G4) and CpG-islands^1, 16, 19, 20^. Interestingly, origins identified by bubble-seq correlate with DNAse hypersensitive regions and the 5’ end but not the body of active transcription units^21^. Both SNS- and bubble-seq detect a higher origin density in early RTDs than in mid-to-late RTDs^19, 21^. Strand-oriented sequencing of Okazaki fragments (OK-seq) reveals the direction of replication forks allowing the mapping of initiation and termination events^22–24^. Bubble-seq^21^ and OK-seq^25^ and DNA combing^26^ studies of mammalian cells demonstrated that replication initiates in broad initiation zones, characterized by flexible initiation from random sites. OK-seq studies identified early initiation zones that are often flanked by actively transcribed genes and are especially enriched in open chromatin, while flanking transcribed gene bodies are enriched in replication termination events^24, 25^. In contrast, in late RTDs, initiation zones are distantly located from active genes and termination occurs over very broad, gene-poor segments.

Comparing replication activation events resulting from SNS-seq, bubble-seq, and OK-seq, the highest concordance was observed between initiation zones detected by OK-seq and bubble-seq^25^.

Chromatin immunoprecipitation followed by next-generation sequencing (ChIP-seq) is a complementary method to map binding sites of origin licensing proteins. ORC and MCM chromatin binding and their relationships with nuclear organization and chromatin features are essential to understand the emergence of replication patterns and replication timing. Drosophila ORC ChIP-seq suggests a stochastic binding pattern often colocalizing at open chromatin marks found at TSS^27^. Genome-wide MCM mapping experiments revealed that this complex is initially loaded at ORC binding sites in absence of Cyclin E/CDK2 activity. With the rise in Cyclin E/CDK2 activity in late G1, MCM is abundantly loaded and redistributed, resulting in a loss of spatial overlap with ORC^13^. In humans, ChIP-seq experiments with single ORC subunits led to the identification of 13,000 to 52,000 potential ORC binding sites^28, 29^. In a previous study, we compared the number of licensed origins in the EBV genome with single replication initiation events and found an excess of 5-10 licensing origins established per genome^30^. A recent genome-wide Mcm7 binding study in human HeLa cells proposed that MCM binds in excess regardless of the chromatin environment, but that origin activation preferentially occurs upstream of active TSSs^31^.

Here, we present the first comparative survey of four different pre-RC components and replication initiation events in the human genome by combining ChIP-seq and OK-seq analyses in the lymphoblastoid Raji cell line. We perform ORC and MCM ChIP in pre-replicative (G1) and post-replicative chromatin, to obtain a comprehensive picture of ORC/ MCM behavior before and after replication. We find ORC and MCM broadly distributed over the genome. In early replicating domains, active transcription locally influences ORC and MCM positioning and consequently replication initiation profiles. In particular MCM are displaced from actively transcribed gene bodies in G1, while ORC is enriched at active TSS. MCM are present at TSS only in post-replicative chromatin. We show that H4K20me3 is present in a subset of non-genic late replicating initiation zones, which are enriched in ORC/ MCM binding. This confirms our previous finding that H4K20me3-mediated ORC-DNA binding enhances origin activity in certain environments^18, 32^. Finally, we find that the global density of ORC highly correlates with replication timing, an effect observed less prominently for MCM. These results argue that ORC but not MCM may dictate replication timing.

## Results

### MODERATE AVERAGING IS THE BEST APPROACH FOR ORC AND MCM DISTRIBUTION ANALYSIS

To obtain a complete picture of ORC and MCM distributions prior to replication initiation, we cell-cycle fractionated human lymphoblastoid Raji cells by centrifugal elutriation into a pre-replicative G1 population (hereafter referred to as *pre*) - which is enriched for ORC/MCM bound chromatin^30^ - and a post-replicative cell population (hereafter referred to as *post*), including S, G2 and mitotic cell populations. Propidium Iodide staining followed by FACS (Supplementary Figure 1a) and Western blot analyses of cyclins A, B, and H3S10 phosphorylation (Supplementary Figure 1b) confirmed the cell cycle stages. To ensure unbiased detection of ORC and MCM positions by ChIP-seq, we simultaneously targeted two members of each complex: Orc2, Orc3, Mcm3 and Mcm7, using validated ChIP-grade antibodies^30, 33, 34^. ChIP efficiency and quality were measured using the Epstein-Barr virus latent origin *oriP* as reference (Supplementary Figure 1c). In EBV, EBNA1 recruits ORC to the *oriP* dyad symmetry element DS. Consequently, we detected ORC at DS in a cell-cycle independent manner, while the presence of MCM was cell-cycle dependent^30, 33^, as expected.

ChIP-sequencing of two ORC (Orc2, Orc3) and of three MCM (Mcm3, Mcm7) replicates in both *pre*- and *post*-fractions resulted in reproducible, but dispersed ChIP-seq signal accumulations at the established replication origin Mcm4/PRKDC (Fig. 1a (*pre*), Supplementary Figure 2a (*post*)). We first employed the MACS2 peak-calling program^35, 36^, but found that the obtained results were too dependent on the chosen settings and concluded that ORC/ MCM distribution was too dispersed to be efficiently captured by peak calling (data not shown), requiring an alternative approach.

**Figure 1:**
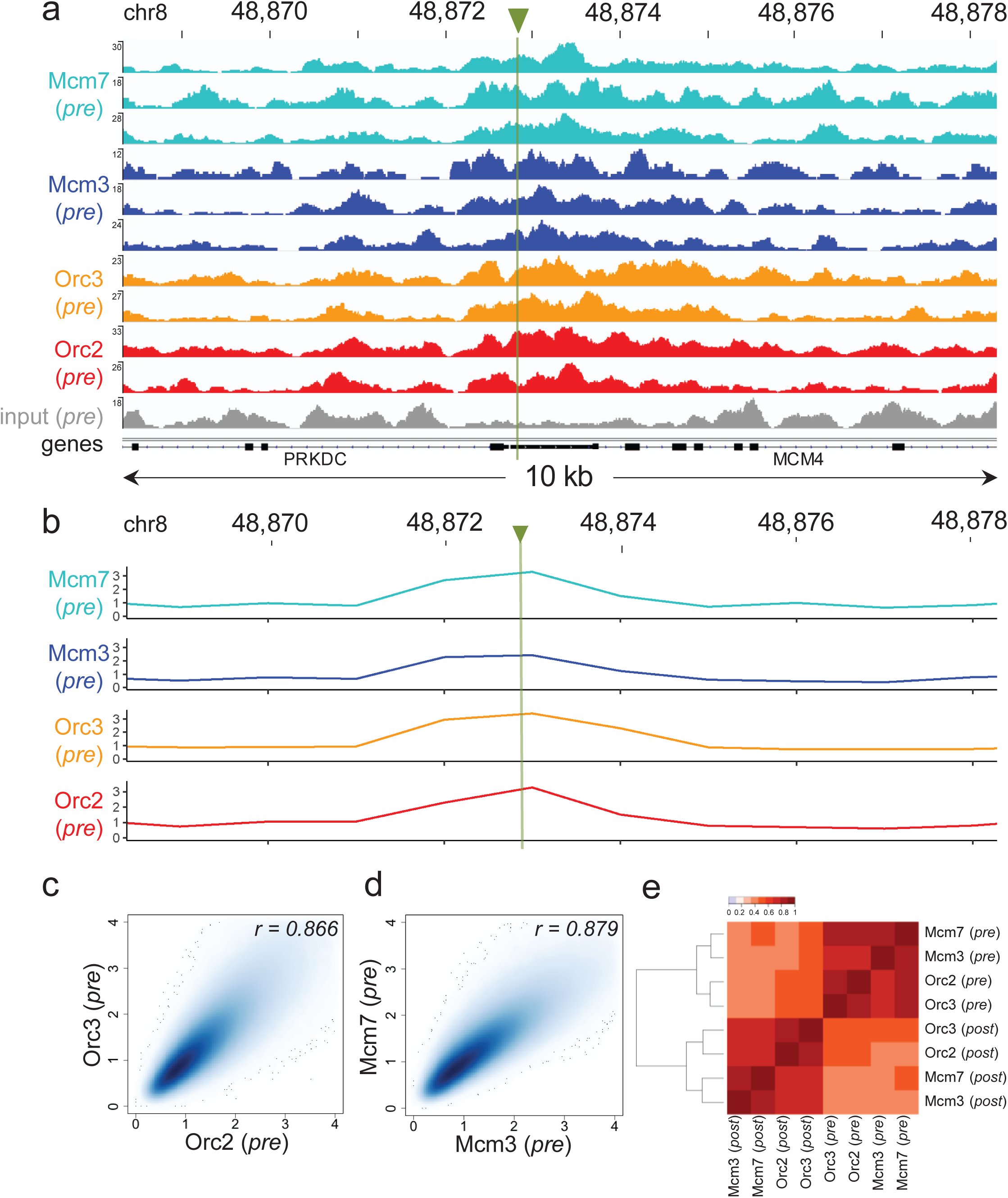
ORC/ MCM ChIP-seq is best analyzed using a moderate averaging approach. a) Sequencing pileup IGB visualization at the Mcm4/PRKDC origin: two samples of Orc2, Orc3 and three samples of Mcm3, Mcm7 (*pre*-fraction), plotted against the input. The profiles are shown in a 10 kb window (chr8: 48,868,314 - 48,878,339), the position of the origin is indicated as green line. Track heights represent raw read depth. b) The profile of ORC/ MCM ChIP-seq after 1 kb binning at the same locus. The reads of replicates were summed and normalized by the total genome-wide ChIP read frequency followed by input division. Y-axis represents the resulting relative read frequency. c) Correlation plot between Orc2 and Orc3 relative read frequencies in 1 kb bins. d) Correlation plot between Mcm3 and Mcm7 relative read frequencies in 1 kb bins. e) Heatmap of Pearson correlation coefficients r between all *pre*- and *post*-fraction ChIP relative read frequencies in 1 kb bins. Column and line order were determined by complete linkage hierarchical clustering.

Consequently, we summed up the reads of the ChIP replicates and combined the number of ChIP-seq reads in 1 kb bins and normalized this signal against the mean read frequency of the entire ChIP sample, followed by input division. We chose 1 kb bins as this window size averages out experimental variations due to the stochastic binding of ORC/ MCM. At the same time, this window is small enough to detect local changes in the binding patterns. As a proof for this assumption, ChIP enrichments at the Mcm4/PRKDC origin were detected after binning (Fig. 1b (*pre*), Supplementary Figure 2b (*post*)). The relative read frequencies of Orc2/Orc3 (Fig. 1c) and Mcm3/Mcm7 (Fig. 1d) showed high Pearson correlation coefficients of r = 0.866 and r = 0.879, respectively. The correlation between ORC and MCM was only slightly lower (Mcm3/Orc2/3: r = 0.775/0.757, Mcm7/Orc2/3: r = 0.821/0.800, Fig. 1e). Hierarchical clustering based on Pearson correlation between ChIP profiles showed that ORC and MCM profiles clustered together, independently from the cell cycle stage. We conclude that this binning approach is valid for analyzing our ChIP-seq data.

Miotto et al. demonstrated that Orc2 positions highly depend on chromatin accessibility and colocalize with DNase hypersensitive (HS) sites present at active promoters or enhancers^29^. Furthermore, Sugimoto et al. observed that active origins, enriched with MCM7 correlate with open chromatin sites^31^. As a DNase HS profile of the Raji cell line does not exist, we compared the ENCODE dataset of DNase HS clusters from 125 cell lines with ORC and MCM read frequencies. We indeed found a significant enrichment of ORC and MCM at DNase HS regions larger than 1 kb, compared to regions deprived of DNase HS sites (Supplementary Figure 3a (*pre*) and 3b (*post*)), validated our data further.

### ORC/MCM ARE ENRICHED IN ZONES OF REPLICATION INITIATION DEPENDENT ON TRANSCRIPTION

After confirming the validity of the ChIP experiments and establishing an analysis approach based on moderate averaging, we compared the relative read frequencies of each pre-RC component to active replication initiation units. Using OK-seq in Raji cells^37^, we calculated the replication fork directionality (RFD), and delineated zones of preferential replication initiation as ascending segments (AS) of the RFD profile. OK-seq does not detect single replication initiation events, but regions of preferential replication initiation (AS)^22, 24, 25^. To assess ChIP signals within AS, we only kept AS of sizes > 20 kb. Using the RFD shift across the AS (ΔRFD) as a measure of replication initiation efficiency, we further required ΔRFD > 0.5 to make sure AS corresponded with efficient initiation zones. In total, we selected 2,957 AS, with an average size of 52.3 kb, which covered 4.9% (155 Mb) of the genome (Fig.2a, green bars, Table 1). 2,451 (83%) of all AS located close to genic regions (AS extended by 20 kb on both sides overlapped with at least one annotated gene). Thereby, 673 AS (22.8% of all AS) were flanked by actively transcribed genes (TPM > 3) at both sides (type 1 AS) with less than 20 kb between AS borders and the closest transcribed gene.1,026 AS (34.7%) had only one border associated to a transcribed gene (type 2 AS). 506 AS (17.1%) were devoid of proximal genes (non-genic AS), where 20 kb extended AS did not overlap with any annotated gene (Table 1). Although the slope did not change considerably in the different AS types, type 1 AS were on average the most efficient, while non-genic AS were slightly less efficient (Supplementary Figure 4a). Furthermore, type 1 and type 2 AS located to early replication timing domains, while non-genic AS were predominantly late replicating (Supplementary Figure 4b), which is in agreement with AS previously described for GM06990 and HeLa^25^.

**Table 1:**
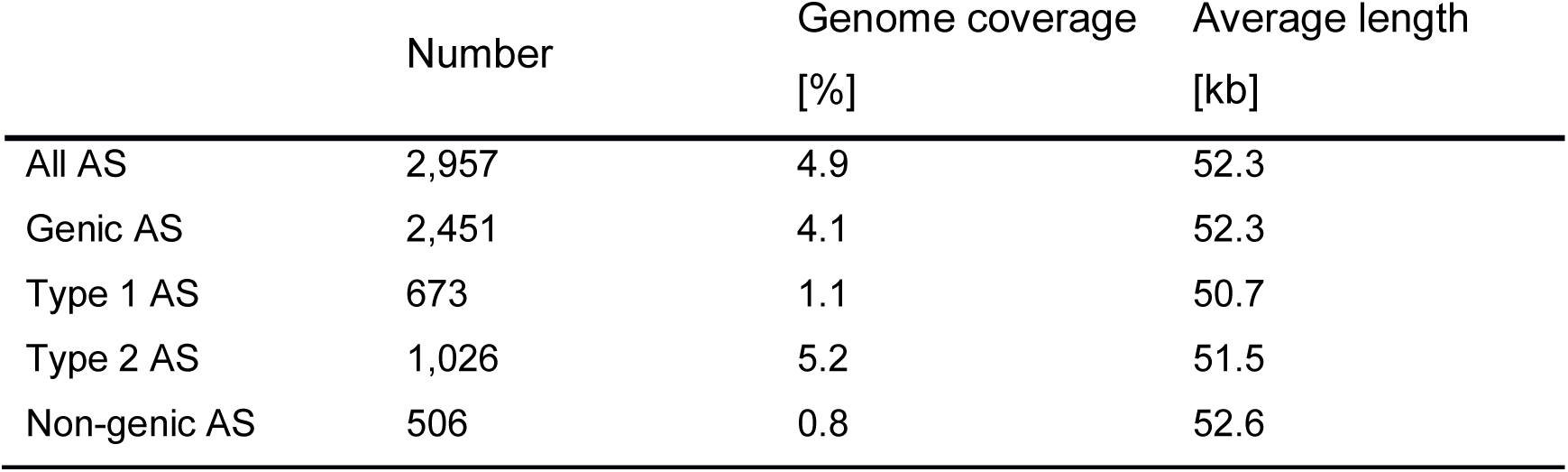
Characterization of different AS subtypes. Only AS ≥ 20kb were considered. Genic AS: flanked by genic region(s) irrespective of transcriptional activity ± 20kb of the AS border. Type 1 and type 2 AS: AS flanked by expressed genes (TPM ≥ 3) within 20kb on both sides (type 1) or one side (type 2). Non-genic: no annotated gene ± 20kb of AS border.

|  | Number | Genome coverage<br>[%] | Average length<br>[kb] |
| --- | --- | --- | --- |
| All AS | 2,957 | 4.9 | 52.3 |
| Genic AS | 2,451 | 4.1 | 52.3 |
| Type 1 AS | 673 | 1.1 | 50.7 |
| Type 2 AS | 1,026 | 5.2 | 51.5 |
| Non-genic AS | 506 | 0.8 | 52.6 |

Replication can only be activated, when functional pre-RCs are established in the preceding G1 phase. We set our ORC/ MCM ChIP-seq signals in relation to RFD and computed the relative read frequencies of ORC/ MCM around all AS aggregate borders. In the *pre-*fraction, both ORC and MCM were enriched within AS compared to flanking regions (Fig. 2b).

**Figure 2:**
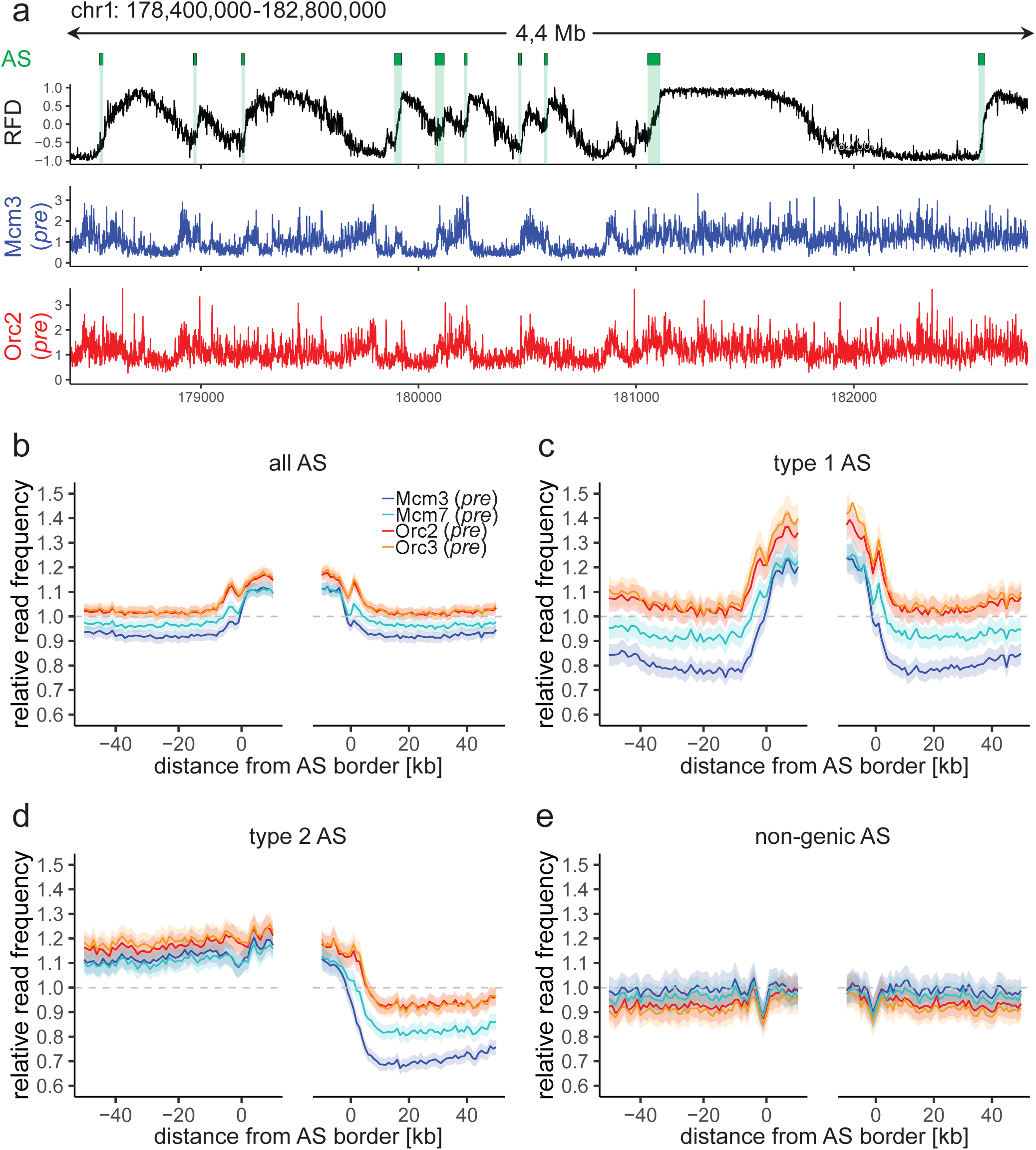
ORC/ MCM enrichment within AS depends on active transcription. a) Top: Example of an RFD profile on chr1: 178,400,000 – 182,800,000. Detected AS are labeled by green rectangles. Bottom: Representative Mcm3 (blue) and Orc2 (red) profile (*pre*-faction) after binning for the same region. b-e) Average relative ChIP read frequencies of Orc2, Orc3, Mcm3, and Mcm7 *pre*-fractions at AS borders of b) all AS, c) type 1 AS with transcribed genes at both AS borders, d) type 2 AS with transcribed genes oriented at their right AS border, and e) non-genic AS in gene deprived regions. The mean of ORC and MCM frequencies are shown ± 2 x SEM (lighter shadows). The dashed grey horizontal line indicates relative read frequency 1.0 for orientation.

To resolve the impact of transcriptional activity, we repeated this calculation and sorted for type 1 AS (Fig. 2c), type 2 AS (Fig. 2d), and non-genic AS (Fig. 2e). This analysis revealed that transcriptional activity in AS flanking regions did not only lead to increased ORC levels inside AS (comparing Fig. 2b and Fig. 2c), but also resulted in a prominent depletion of MCM from transcribed flanking regions (Figs. 2c and 2d). In contrast, in type 2 AS, ORC/ MCM levels remained elevated at AS borders without transcriptional activity (Fig. 2d, left border), and no evident ORC/ MCM enrichments were detected within non-genic AS (Fig. 2e). *Post*-fraction profiles display the same tendencies, however to a lesser extent (Supplementary Figure 5a-d), implying a partial displacement of ORC/ MCM during S-phase. AS borders characterized by transcriptional activity were locally enriched by ORC and MCM. This is in line with previously detected Orc1 accumulation at AS borders^25^ and indicates that the *post-*fraction contains an important portion of cells in late mitosis, when origin licensing is initiated.

### ORC IS ENRICHED AT TSS OF ACTIVELY TRANSCRIBED GENES AND MCM DEPLETED FROM GENE BODIES

Replication initiation often correlates with active gene transcription^16, 19, 38, 39^. A recent study using OK-seq even linked both replication initiation and termination to transcription^24^. Furthermore, ORC/MCM enrichment in type 1 and 2 AS compared to genic flanking regions (Fig. 2c and d) argue for a major contribution of active transcription to ORC/ MCM positioning. To study the association of ORC/ MCM localizations and transcriptional activity, we set our ChIP-seq data in relation with transcription profiles obtained from asynchronously cycling Raji cells. We analyzed ORC/ MCM relative read frequencies around active TSS and transcriptional termination sites (TTS) (Fig. 3). ORC relative read distribution of G1-phased cells (*pre*) was significantly enriched at active TSS as already demonstrated in *Drosophila*^27^, and human cells^28, 29^. Relative read frequency levels of ORC were moderately but significantly higher upstream of TSS and downstream of TTS than within genes (Fig. 3a). Approximately 45% of actively transcribed gene bodies were significantly depleted from ORC in the *pre*-fraction (Supplementary Table 1).

**Figure 3:**
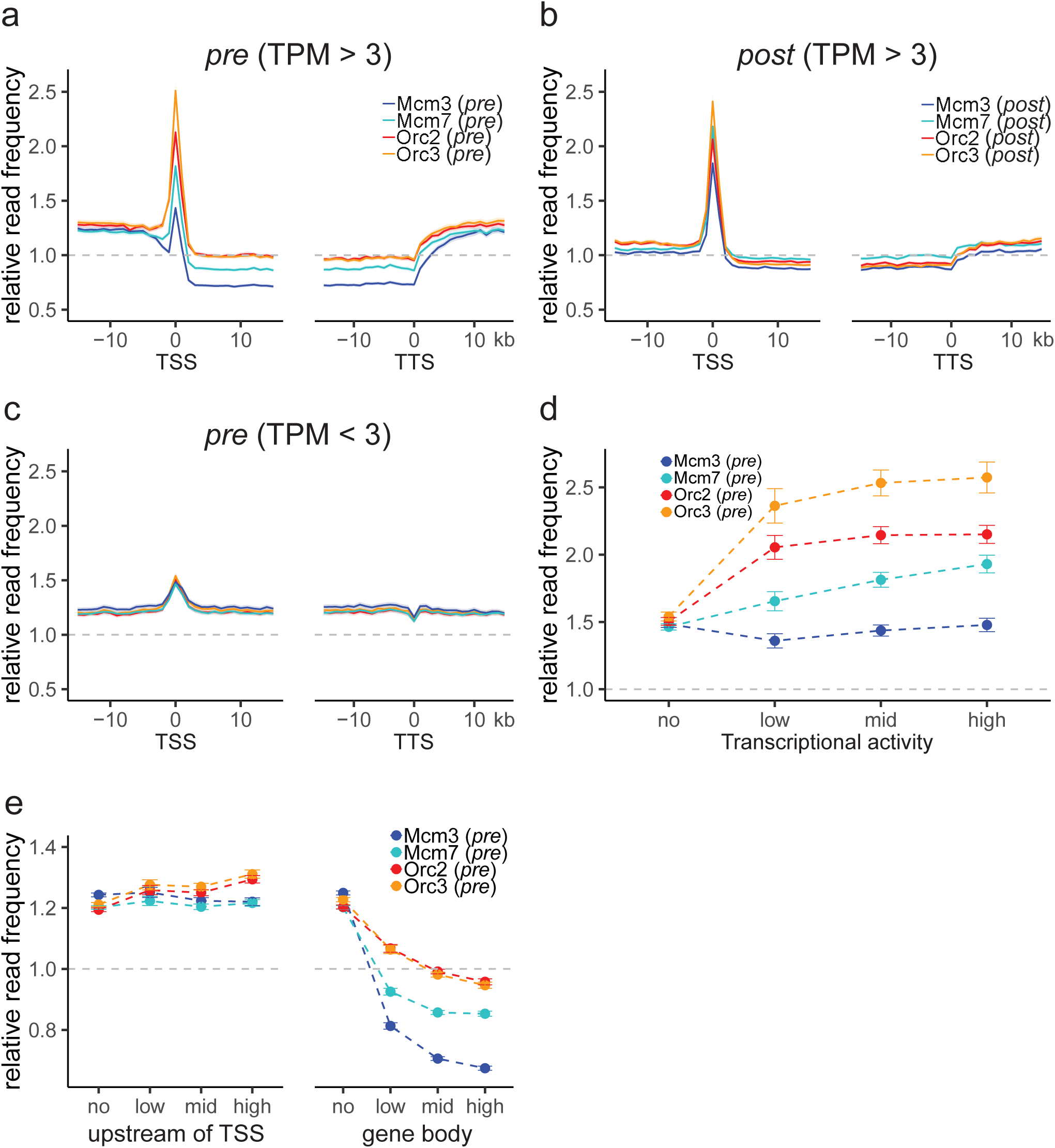
ORC is enriched at active TSS while especially MCM is depleted from actively transcribed genes. a) - c) Relative ORC/ MCM read frequencies around active TSS or TTS for a) active genes (TPM > 3) in *pre*, b) active genes (TPM > 3) in *post*, and c) inactive genes (TPM < 3; *pre*-fraction). Only genes larger than 30 kb without any adjacent gene within 15 kb were considered. Distances from TSS or TTS are indicated in kb. Means of ORC and MCM frequencies are shown ± 2 x SEM (lighter shadows). d) ORC/ MCM (*pre*) read frequencies at TSS dependent on transcriptional activity (± 2 x SEM). e) ORC/ MCM (*pre*) relative read frequencies upstream of TSS and in the gene body dependent on transcriptional activity (± 2 x SEM; TSS ± 3 kb removed from analysis). Transcriptional activity was classified as: no (TPM < 3), low (TPM 3-10), mid (TPM 10-40), high (TPM > 40). The dashed grey horizontal line indicates relative read frequency 1.0 for orientation.

Compared to ORC, MCM enrichment at TSS was less prominent, however, depletion from gene bodies was more pronounced (Fig. 3a). 75% and 58% of investigated transcribed gene bodies were significantly depleted from Mcm3 and Mcm7, respectively (Supplementary Table 1). Interestingly, while ORC profiles did only slightly change from *pre*- to *post*-fractions (Fig. 3a vs. Fig. 3b), MCM profiles of the *post*-fraction rather resembled ORC profiles, with a significant peak at TSS and a less pronounced depletion from gene bodies (Fig. 3b). This observation is explained by a drastic reduction of the number of significant MCM depleted genes in the *post*-fraction (from 75.2% (Mcm3) and 58.3% (Mcm7) in *pre*-fractions to 28.5% and 16.7% in *post*-fractions). The number of ORC depleted genes only decreased from 44% in *pre*- to 34-38% in *post*-fractions (Supplementary Table 1).

Chen *et al*. recently reported that transcriptional activity models the replication initiation profile^24^. We found that TSSs of inactive genes were hardly enriched for ORC/ MCM and that inactive gene bodies were not depleted from licensing components (Fig. 3c). Although active transcription is necessary for ORC enrichment at TSS, we also observed that increasing transcriptional activity did not have any major impact on ORC/ MCM enrichments at TSS (Fig. 3d, Supplementary Fig. 6a (*post*)). The same is true for ORC/ MCM depletion from gene bodies in *pre*- or *post-* fractions (Fig. 3e and Supplementary Fig. 6b).

The *pre*-fraction represents a cell cycle stage immediately prior to origin activation, with an excess of MCM loaded onto chromatin^13, 40^. Here, we found MCM being actively displaced from gene bodies by the transcriptional machinery, as previously proposed in *Drosophila* by Powell *et al*^13^. In post-replicative chromatin, obtained from a cell population containing a prominent fraction of mitotic cells (Supplementary Figure 1b), we found that MCM co-localized with ORC at TSS, possibly reflecting early MCM loading. These findings suggest, that the co-directionality between DNA replication and transcription of active genes is achieved by enhancing pre-RC formation at TSS^24^. Inhibiting origin licensing within active genes contributes to genome stability by preventing intragenic replication initiation and thus colliding events that originate from head-on oriented DNA replication and transcription^41^.

### LATE REPLICATING AS ARE CHARACTERIZED BY H4K20ME3

In the preceding sections, we showed that the enrichment of ORC and MCM at type 1 and type 2 AS depended on transcriptional activity. Non-genic AS were characterized by the absence of any transcriptional annotation. Interestingly, we did not detect a significant accumulation of ORC/ MCM at late replicating non-genic AS (Fig. 2e). Therefore, we asked for other characteristics determining their replication. We recently demonstrated that H4K20me3 supports the licensing of a subset of late replicating origins in heterochromatin^18^ and hypothesized that H4K20me3 may also influence licensing of non-genic AS. We performed ChIP for H4K20me3 and its precursor H4K20me1 in three replicates in *pre*-fractions and validated its success by qPCR (Supplementary Figure 7 a (H4K20me3) and 7b (H4K20me1)). After sequencing, we performed MACS2 broad peak-detection and kept only peaks overlapping in all three samples (16852 peaks for H4K20me3 and 12264 peaks for H4K20me1, see also Supplementary Table 2 for further characterization). H4K20me3 peak sizes ranged from 200 bp to 105 kb (200 bp to 183 kb for H4K20me1, Supplementary Table 2, Supplementary Figure 7c). When calculating ORC/ MCM coverage of the *pre-*fraction at H4K20me3/me1 peaks > 1 kb (12251/ 6277 peaks, respectively), we predominantly detected ORC and also some MCM enrichment at H4K20me3 sites (Fig. 4a, Supplementary Fig. 7d (*post*)). By contrast, H4K20me1 peaks were not enriched in both ORC and MCM (Supplementary Figure 7e).

**Figure 4:**
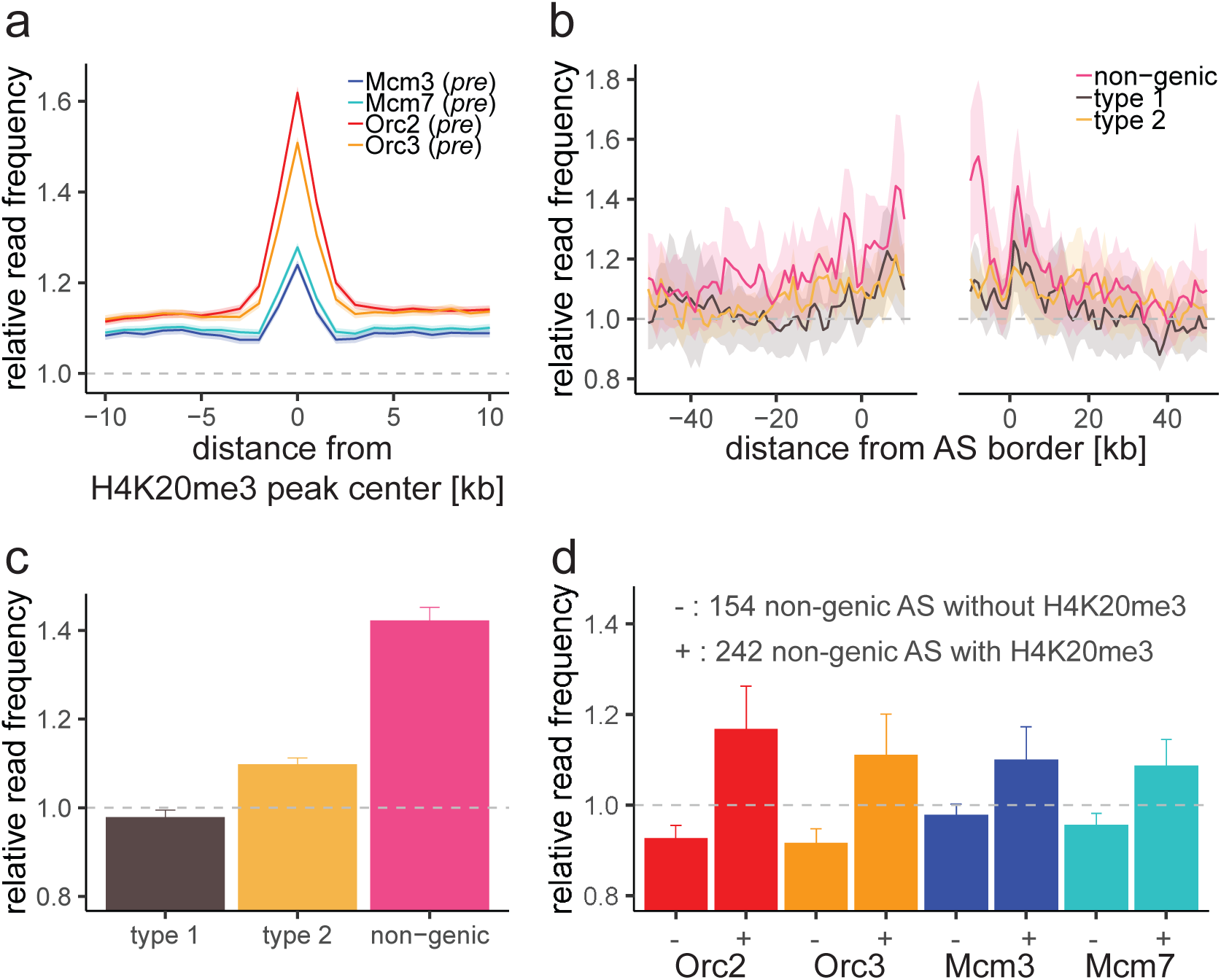
Non-genic late replicating AS containing H4K20me3 are preferentially enriched in ORC/ MCM. a) Average ORC/ MCM relative read frequencies (*pre*-fraction) at H4K20me3 peaks (> 1 kb). b) Cumulative relative ChIP read frequencies of H4K20me3 at AS borders of the different AS types. Means of ChIP read frequencies are shown ± 2 x SEM (lighter shadows). c) Histogram representation of mean H4K20me3 read frequencies ± 2 x SEM within the different AS types. d) Histogram representation of mean ORC/ MCM read frequencies at non-genic AS without (242 non-genic AS) or with (154 non-genic AS) H4K20me3 ± 2 x SEM. The dashed grey horizontal line indicates relative read frequency 1.0 for orientation.

Consequently, we asked, if H4K20me3 was enriched in AS. When calculating H4K20me3 coverage at the different AS types, we specifically detected H4K20me3 in non-genic AS, representing the first histone modification characterizing late replicating AS (Fig. 4b and c). Starting from 506 non-genic AS, we extracted a subset of 154 non-genic AS associated to H4K20me3 (where H4K20me3 relative read frequency was above the genome mean value by more than 1.5 standard deviation), versus 242 non-genic AS with H4K20me3 levels lower than genome average. We found ORC and MCM present at the H4K20me3-associated subgroup compared to non-genic AS without H4K20me3 (Fig. 4d). These results indicate that transcriptionally independent non-genic AS are potentially characterized by specific histone modifications that lead to ORC/ MCM recruitment, albeit remaining undetectable when considering all non-genic AS.

### EARLY REPLICATION TIMING DOMAINS ARE ENRICHED FOR ORC

When assessing ORC/ MCM distributions at a local level, ORC/ MCM seemed to be nearly equally distributed throughout the entire genome with the exception of genic regions. Consequently, we asked whether ORC/ MCM distributions impact on the more global event of replication timing as has been observed in *S. cerevisiae* for MCM^42^. We extracted early and late RTDs from Raji cells using Early/Late Repli-seq data from Sima *et al.*^43^. Employing a threshold of early to late ratio > 1.6 for early RTDs and < -2.0 for late RTDs resulted in 302 early RTDs covering 642.8 Mb and 287 late RTDs covering 617.4 Mb of the genome. Working in 10 kb bins, we removed all bins containing annotated genes in a ± 10 kb window from the analysis, to obtain data independent from transcriptional activity. Calculating the mean ORC/ MCM relative read frequencies of the *pre*-fraction in early compared to late RTDs revealed ORC being 1.4-times enriched in early RTDs compared to late RTDs (Fig. 5a; Supplementary Table 3). By contrast, MCM were less enriched in early and less depleted from late RTDs (Fig. 5a; Supplementary Table 3). In the *post*-fraction, ORC enrichment reduced to a ratio early/late of 1.2, while especially Mcm3 relative read frequencies were smaller in early RTDs than in late RTDs (Fig. 5b; Supplementary Table 3). We consequently conclude, that elevated ORC levels are tightly associated with early replication timing throughout the cell cycle, while ORC levels decrease below average in late RTDs. MCM levels however, albeit showing the same tendencies, seem to be less assertive.

**Figure 5:**
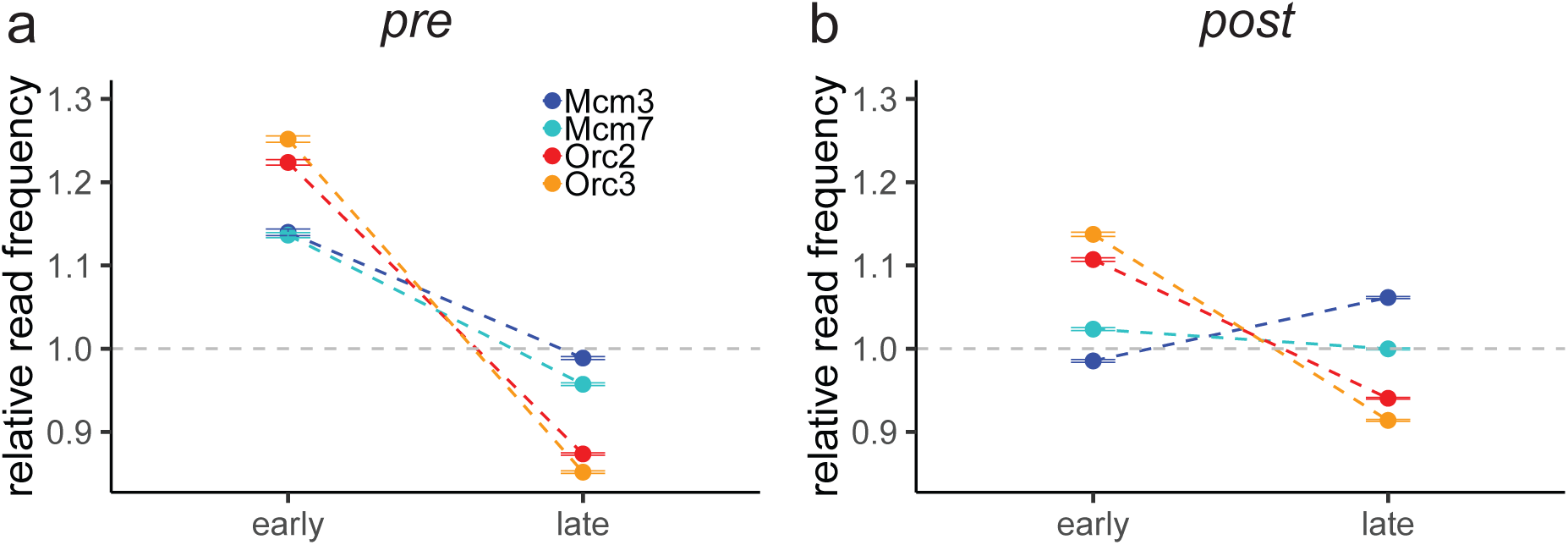
ORC is highly enriched in early RTDs. a-b) Mean ORC/ MCM read frequencies (± 2 x SEM) in early or late RTDs of a) the *pre*-fraction or b) the *post*-fraction. The analysis was performed in 10 kb bins. Any gene ± 10 kb was removed from the analysis. The dashed grey horizontal line indicates relative read frequency 1.0 for orientation.

In summary, we demonstrate that ORC abundance correlates with early replication timing and propose a model in which ORC/ MCM positioning is strongly affected by transcriptional activity in early RTDs (Fig. 6a). However, the association of origin licensing and replication initiation in late RTDs with specific histone modifications, e.g. H4K20me3 (Fig. 6b), illustrates that ORC/ MCM alone are not sufficient to explain the observed replication initiation profiles. Missing factors likely remain to be described e.g. at the transcription free border of type 2 AS.

**Figure 6:**
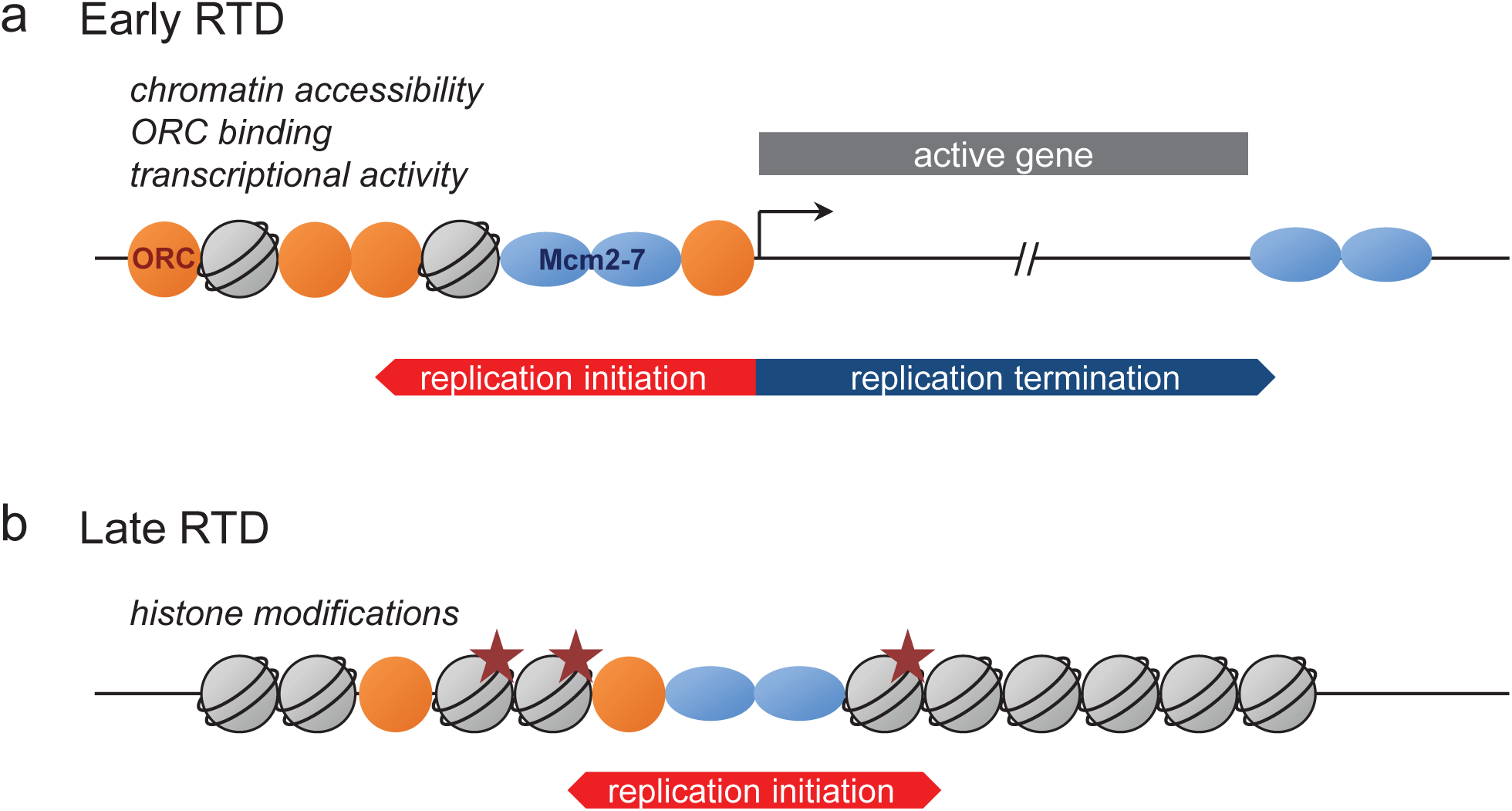
Model of replication regulation in early and late RTDs. a) In transcriptionally active, early RTDs, ORC preferentially binds active TSS, where it also loads MCM, which is actively displaced by transcription. The combination of chromatin accessibility, ORC binding and transcriptional activity define replication initiation and termination zones. b) In gene deprived, late RTDs, ORC specifically binds to histone modifications, such as H4K20me3. Lower levels of MCM loading are sufficient for replication initiation.

## Discussion

The study presented here provides a novel comprehensive genome-wide analysis of multiple pre-RC proteins and replication initiation in human cells. We find that on the local level, ORC and MCM are enriched in zones of replication initiation, especially in early replicating domains. Active transcription highly influences the distribution of licensed origins: ORC accumulates at active TSS and especially MCM is depleted from actively transcribed gene bodies (Fig. 3). We demonstrate that late replicating non-genic AS associated with H4K20me3 are characterized by elevated ORC/ MCM levels. When looking at the global level of replication timing, we find that high ORC and, to a lesser extent MCM, correlate with early replication timing, whereas late RTDs are deprived of both ORC and MCM (Fig. 5).

### TRANSCRIPTIONAL ACTIVITY STRONGLY INFLUENCES LOCAL ORC/ MCM DISTRIBUTION

Type 1 and type 2 AS are at their transcribed borders are characterized by drastic changes in the occupancy of ORC and MCM (Fig. 2). More specific, actively transcribed gene bodies were devoid of MCM, while ORC and, to a lesser degree, MCM were detected at active TSS (Fig. 3a). Remarkably, a prominent peak at TSS was also observed for MCM in *post*-chromatin (Fig. 3b). Thereby, the prominence of this peak strictly correlates with transcriptional activity of the neighboring gene, although not with the transcriptional level (Fig. 3d, Supplementary Fig. 6a). These findings suggest that active TSS are efficient in replication licensing. Two possibilities might explain this feature: First, TSS are hotspots of ORC binding as they are easily accessible and represent sites where MCM re-association starts for the next cell cycle when CDK activity is low^44^. In the following G1 phase, MCM complexes are distributed from these sites to upstream regions but not downstream into transcribed gene bodies. Depletion of ORC and MCM from actively transcribed gene bodies is independent from the level of transcriptional activity (Fig. 3e and Supplementary Fig. 6b). The displacement of MCM might be due to the moving RNA polymerase II, as suggested by Powel et al.^13^. Second, residual amounts of MCM are being observed in MCM-immunoblots of chromatin association experiments in G2/M^33^. These MCM molecules might be located at TSS and remain on chromatin throughout the cell cycle. The prominent peak is only visible, if the majority of intergenic MCM has been displaced from chromatin after replication during S phase. In this hypothesis, TSS would constitute cell cycle-independent MCM storage sites.

Initiating DNA replication at TSS reduces the risk colliding replication and transcription forks^41^.

### LATE REPLICATING NON-GENIC AS CORRELATE WITH H4K20ME3 BUT MAY REQUIRE ADDITIONAL FEATURES

H4K20 methylation has multiple functions in ensuring genome integrity, such as DNA replication^39, 45, 46^, DNA damage repair, and chromatin compaction^32, 47, 48^ suggesting that the different functions are context dependent and executed with different players. We previously demonstrated that H4K20me3 provides a platform to enhance origin formation in late replicating heterochromatin^18^. Shoaib and colleagues reported recently that H4K20me3 restricts replication licensing to prevent overreplication^32^. The latter is in line with our observation that mainly ORC but little MCM are enriched at H4K20me3 peaks (Fig. 4a). However, when selecting for H4K20me3-containing non-genic AS, we detect both elevated ORC and MCM in this particular AS subset, suggesting that the amount of MCM is sufficient for replication activation. Yet, we could not detect a general ORC/ MCM enrichment in all non-genic AS, and it is currently unclear which additional features may be required for specifying or activating non-genic AS. Together with our previous observations^18^, we conclude that H4K20me3 is pivotal for the replication of at least a subset of transcriptionally silent non-genic AS, although there might be still unknown factors associated to other subsets of non-genic AS.

### DISPERSED MCM BINDING IS CONSISTENT WITH THE CASCADE MODEL

While the genome-wide distributions of ORC and MCM are convincingly mapped by high-throughput ChIP in *S. cerevisiae*^49^ and in Drosophila^13, 27^, the analysis of pre-RC chromatin binding in human cells still remains a special challenge. To date, only few genome-wide studies exist for human ORC and Mcm7, which are identifying binding sites located in early replicating regions near transcribed genes^28, 29, 31, 50^. Replication of metazoan genomes is organized into subnuclear domains, each containing several clusters^5, 6, 51^. Within a cluster, replication initiates stochastically from an excess of dispersed licensed origins. This leads, in comparison to site-specific chromatin binding factors^52^, to low enrichments of ORC in ChIP experiments and in particular of MCM, which spreads after chromatin loading^14^. By choosing a moderate cumulative approach, we were able to eliminate variations due to stochastic ORC/ MCM binding, while conserving local changes in their binding patterns.

Our data is in line with the previously proposed cascade model that replication of the human genome involves a superposition of efficient initiation at “master” origins identified by RFD AS, followed by a cascade of disperse, less efficient origin activation along the intervening domain^53^. We found a clear depletion of ORC and MCM in transcription units that border type 1/2 AS, but no convincing depletion of MCM in the rest of the non-transcribed genome (e.g. compare type 2 AS with non-transcribed flank (Fig 2d, left side) and non-genic AS with both flanks (Fig. 2e)). While these data favor a broad dispersion of potential origins within and outside “master” initiation zones, consistent with the cascade model, we also suggest that a higher MCM density *is not* a distinguishing feature of AS from the rest of the (non-transcribed) genome - except perhaps for a subset of non-genic AS associated with H4K20me3. Instead, initiation zone specification may occur at the origin activation rather than at the licensing step and may be mediated by preferential accessibility to limiting factors during S phase^5^. Increased accessibility of MCMs within AS might be related to the co-enrichment for several active chromatin marks specifically found in initiation zones even when they are not flanked by active genes, e.g. DNAse HS sites, H3K4me1, H3K27ac, p300^54^.

### GLOBAL ORC DISTRIBUTION CORRELATES WITH REPLICATION TIMING

The spatio-temporal replication program is relatively well conserved in consecutive replication cycles for each cell type, differs only slightly between cell lines and changes during differentiation^55^. Many studies have shown that the timing program correlates with topological domains and all origins within one domain replicate in the same time frame^3, 51^. Topological domains are remarkably stable, explaining why the spatial replication profile is conserved^56^. Early RTDs correlate with topological domains enriched in ORC and are characterized by active transcription and chromatin activating features, as previously shown^24^ and confirmed by our data. Currently it is controversially discussed, if higher amounts of ORC^28, 29^ or the excess of reiteratively loaded MCM^14, 42^ determine replication timing.

Our results imply that the global density of ORC correlates with replication timing. This observation is independent from local transcriptional influences, as we removed all genes including 10kb of flanking regions from our analyses. We propose that highly dynamic euchromatin generates advantageous conditions allowing the binding of ORC to chromatin (Fig. 6a). The spatial association of MCM to chromatin is actively regulated by transcription as transcribed gene bodies are kept clear of licensing (Fig. 6a). The organizational link between active transcription and replication directly ties transcriptional programs and replication patterns during cellular re-organization, as for example differentiation. Comparing replication licensing and initiation patterns in pluripotent stem cells and differentiated cells will give further insights in this functional connection of transcription and replication.

Heterochromatin is predominantly replicated late in S phase and ORC binding is enhanced through specific interactions with histone modifications (e.g. H4K20me3^18^). This leads to lower global ORC levels in late RTDs, compared to early RTDs (Fig. 5a). However, we speculate that these specific ORC-chromatin interactions lead to sufficient MCM loading (Fig. 6b), as we observe higher MCM levels than ORC levels in late RTDs.

The flexibility in origin activation and the stochastic use of replication origins allows the cell to adapt to environmental constraints. In summary, we found origin licensing throughout the genome, which allows the cell to activate replication wherever needed. In early replicating chromatin, the origin licensing and initiation pattern is tightly connected with the transcriptional program, whereas in late replication domains other factors, including H4K20me3, are shaping these replication processes.

## Online Methods

### Cell culture

Raji cells (ATCC) were cultivated at 37°C and 5% CO2 in RPMI 1640 (Gibco, Thermo Fisher, USA) supplemented with 8% FCS (Lot BS225160.5, Bio&SELL, Germany), 100 Units/ml Penicillin/ 100 µg/ml Streptomycin (Gibco, Thermo Fisher, USA), 1x MEM non-essential amino acids (Gibco, Thermo Fisher, USA), 2 mM L-Glutamin (Gibco, Thermo Fisher, USA), and 1 mM Sodium pyruvate (Gibco, Thermo Fisher, USA).

### RNA extraction, sequencing, TPM calculation

RNA was extracted from 3 x 10^5^ Raji cells using Direct-zolTM RNA MiniPrep kit (Zymo Research) according to manufacturers’ instructions. RNA quality was confirmed by Bioanalyzer RNA integrity numbers between 9.8 and 10 followed by library preparation (Encore Complete RNA-Seq Library Systems kit (NuGEN)). Single-end 100 bp sequencing was performed by Illumina HiSeq 1500 to a sequencing depth of 25 million reads. The reads were mapped to hg19 genome using Tophat2 and assigned to annotated genes (HTSeq-count)^57, 58^. TPM values (Transcripts per kilobase per million reads) were calculated for each sample 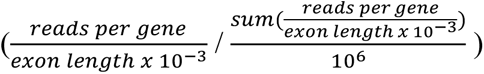 as previously described^59^.

### Replication fork directionality profiling using OK-seq method in Raji^37^

Raji OK-seq was recently published as part of Wu *et al*. and is available from the European Nucleotide Archive under accession number PRJEB25180 (see data access section)^37^. Reads > 10 nt were aligned to the human reference genome (hg19) using the BWA (version 0.7.4) software with default parameters^60^. We considered uniquely mapped reads only and counted identical alignments (same site and strand) as one to remove PCR duplicate reads. Five biological replicates were sequenced providing a total number of 193.1 million filtered reads (between 19.1 and 114.1 million reads per replicate). RFD was computed as 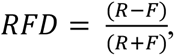 where “R” (resp. “F”) is the number of reads mapped to the reverse (resp. forward) strand of the considered regions. RFD profiles from biological replicates were highly correlated, with Pearson correlation computed in 50 kb non-overlapping windows with > 100 mapped reads (R+F) ranging from 0.962 to 0.993. Reads from the 5 replicate experiments were pooled together for further analyses.

### Determining regions of ascending and descending RFD

RFD profiling of 2 human cell lines revealed that replication primarily initiates stochastically within broad (up to 150kb) zones and terminates dispersedly between them. These initiation zones correspond to quasi-linear ascending segments (AS) of varying size and slope within the RFD profiles. As previously described for mean replication timing profiles analysis^61, 62^, we determined the smoothed RFD profile convexity from the convolution with the second derivative of the Gaussian function of standard deviation 32 kb. 4891 AS were delineated as the regions between positive and negative convexity extrema of large amplitude. The amplitude threshold was set in a conservative manner in order to mainly detect the most prominent initiation zones described by Petryk et al. and avoid false positives^25^. Noting *pos_5’* and *pos_3’* the location of the start and end position of an AS, each AS was associated to its size *pos_3’-pos_5’* and the RFD shift across its length: ΔRFD = RFD (*pos_5’*) – RFD (*pos_3’*).

### Centrifugal elutriation and flow cytometry

For centrifugal elutriation, 5 x 10^9^ exponentially growing Raji cells were harvested, washed with PBS and resuspended in 50 ml RPMI 1680/ 8% FCS/ 1mM EDTA/ 0.25 U/ml DNaseI (Roche, Germany). Concentrated cell suspension was passed through 40 µm cell strainer and injected in a Beckman JE-5.0 rotor with a large separation chamber turning at 1500 rpm and a flow rate of 30 ml/min controlled by a Cole-Parmer Masterflex pump. While rotor speed was kept constant, 400 ml fractions were collected at increasing flow rates (40, 45, 50, 60, 80 ml/min). Individual fractions were quantified, 5 x 10^6^ cells washed in PBS, Ethanol fixed, RNase treated and stained with 0.5 mg/ml Propidium Iodide. DNA content was measured using the FL2 channel of FACSCalibur^TM^ (BD Biosciences, Germany). Remaining cells were subjected to cross-link.

### Cross-link

Raji cells were washed twice with PBS, resuspended in PBS to a concentration of 2 x 10^7^ cells/ml and passed through 100 µm cell strainer (Corning Inc., USA). Fixation for 5 min at room temperature was performed by adding an equal volume of PBS 2% methanol-free formaldehyde (Thermo Scientific, USA) and stopped by the addition of glycine (125 mM final concentration). After washing once with PBS and once with PBS 0.5% NP-40, cells were resuspended in PBS containing 10% glycerol, pelleted and snap frozen in liquid nitrogen.

### Cyclin Western Blot

Cross-linked samples were thawed on ice, resuspended in LB3+ sonication buffer containing protease inhibitor and 10 mM MG132. After sonicating 3 x 5 min (30 sec on, 30 sec off) using Bioruptor in presence of 212-300 µm glass beads, samples were treated with 50 U Benzonase for 15 min at room temperature and centrifuged 15 min at maximum speed. 50 µg protein lysates were loaded on 10% SDS-polyacrylamid gel (Cyclin A1/A2, Cyclin B1), or 12.5%-15% gradient gel (H3S10P). Cyclin A1/A2 (Abcam, ab185619), Cyclin B1 (Abcam, ab72), H3S10P (Cell signaling, D2C8) antibodies were used in 1:1000 dilutions, GAPDH (clone GAPDH3 10F4, rat IgG2c; Monoclonal Antibody Core Facility, Helmholtz Center München) was diluted 1:50. HRP-coupled secondary antibodies were used in 1:10000 dilutions. Detection was done using ECL on CEA Blue Sensitive X-ray films.

### Chromatin sonication

Cross-linked cell pellets were thawed on ice, resuspended LB3+ buffer (25 mM HEPES (pH 7.5), 140 mM NaCl, 1 mM EDTA, 0.5 mM EGTA, 0.5% Sarcosyl, 0.1% DOC, 0.5% Triton-X-100, 1X protease inhibitor complete (Roche, Germany)) to a final concentration of 2 x 10^7^ cells/ml. Sonication was performed in AFA Fiber & Cap tubes (12x12 mm, Covaris, Great Britain) at an average temperature of 5°C at 100W, 150 cycles/burst, 10% duty cycle, 20 min (post fraction: 17 min) using the Covaris S220 (Covaris Inc., UK).

### Chromatin immunoprecipitation and qPCR quality control

Sheared chromatin was pre-cleared with 50 µl protein A Sepharose 4 Fast Flow beads (GE Healthcare, Germany) per 500 µg chromatin for 2h. 500 µg chromatin (or 250 µg for histone methylation) were incubated with rabbit anti-Orc2, anti-Orc3, anti-Mcm3, anti-Mcm7^30^, mouse anti-H4K20me1 (Diagenode, MAb-147-100), rabbit anti-H4K20me3 (Diagenode, MAb-057-050), or IgG isotype control for 16h at 4°C. BSA-blocked protein A beads (0.5 mg/ml BSA, 30 µg/ml salmon sperm, 1X protease inhibitor complete, 0.1% Triton-X-100 in LB3(-) buffer (without detergents)) were added (50 µl/ 500 µg chromatin) and incubated for at least 4h on orbital shaker at 4°C. Sequential washing steps with RIPA (0.1% SDS, 0.5% DOC, 1% NP-40, 50 mM Tris (pH 8.0), 1 mM EDTA) 150mM NaCl, RIPA-300 mM NaCl, RIPA-250 mM LiCl buffer, and twice in TE (pH 8.0) buffer were performed. Immunoprecipitated chromatin fragments were eluted from the beads by shaking twice at 1200 rpm for 10 min at 65°C in 100µl TE 1% SDS. The elution was treated with 80 µg RNAse A for 2h at 37°C and with 8 µg proteinase K at 65°C for 16h. DNA was purified using the NucleoSpin Extract II Kit. Quantitative PCR analysis of the EBV *oriP* DS element (for pre-RC ChIP), or H4K20me1 and -me3 positive loci were performed using the SYBR Green I Master Mix (Roche) and the Roche LightCycler 480 System. Oligo sequences for qPCR were DS_fw: AGTTCACTGCCCGCTCCT, DS_rv: CAGGATTCCACGAGGGTAGT, H4K20me1positive_fw: ATGCCTTCTTGCCTCTTGTC, H4K20me1positive_rv: AGTTAAAAGCAGCCCTGGTG, H4K20me3positive_fw: TCTGAGCAGGGTTGCAAGTAC, H4K20me3positive_rv: AAGGAAATGATGCCCAGCTG. Chromatin sizes were verified by loading 1-2 µg chromatin on a 1.5% agarose gel. Samples were quantified using Qubit HS dsDNA.

### ChIP-sample sequencing

ChIP sample library preparations from > 4 ng of ChIP-DNA was performed using Accel-NGS® 1S Plus DNA Library Kit for Illumina (Swift Biosciences). 50 bp single-end sequencing was done with the Illumina HiSEQ 1500 sequencer to a sequencing depth of ∼ 70 million reads. Fastq-files were mapped against the human genome (hg19, GRCh37, version 2009), extended for the EBV genome (NC007605) using bowtie (v1.1.1)^63^. Pileup profiles were generated in R by extending 50 bp reads by 75 bp at both sites, and calculating the number of reads per base. Sequencing pileup profiles were visualized in Integrated Genome Browser^64^.

For H4K20me1 and -me3 ChIP-seq data in *pre*-fractioned cells, MACS2 peak-calling was performed using the broad setting and overlapping peaks in three replicated were retained for further analyses.

### Binning approach and normalization

All data processing and analysis steps were performed in R (v.3.2.3), visualizations were done using the ggplot2 (v3.1.0) package^65^. The numbers of reads were calculated in non-overlapping 1 or 10 kb bins and saved in bed files for further analysis. To combine replicates, their sum per bin was calculated (= read frequency). To adjust for sequencing depth, the mean frequency per bin was calculated for the whole sequenced genome and all bins’ counts were divided by this mean value. To account for variations in the input sample, we additionally divided by the relative read frequency of the input. This resulted in relative read frequency ranging from 0 to ∼30 Pair-wise Pearson correlations of each ORC/ MCM sample in *pre-* and *post-*fractions were clustered by hierarchical clustering using complete linkage clustering.

### Relation of ChIP relative read frequencies to DNase hypersensitivity

The ENCODE ‘DNase clusters’ track wgEncodeRegDnaseClusteredV3.bed.gz (03.12.2017) containing DNase hypersensitive sites from 125 cell lines were retrieved from http://hgdownload.soe.ucsc.edu/goldenPath/hg19/encodeDCC/wgEncodeRegDnaseCl ustered/. Bins overlapping or not with HS sites larger than 1 kb were defined and the respective ChIP read frequency assigned for comparison.

### Comparison of ChIP relative read frequencies to replication data

AS were aligned on their left (5’) and right (3’) borders. Mean and standard error of the mean (SEM) of relative read frequencies of aligned 1 kb bins were then computed to assess the average ChIP signal around the considered AS borders 50 kb away from the AS to 10 kb within the AS. To make sure bins within the AS were closer to the considered AS border than to the opposite border, only AS of size >20 kb were used (3247/4891). We also limited this analysis to AS corresponding to efficient initiation zones by requiring ΔRFD > 0.5, filtering out a further 290 lowly efficient AS. In order to question the relationship between AS and transcription, we compared the results obtained for different AS groups: 506 AS were classified as non-genic AS when the AS locus extended 20-kb at both ends did not overlap any annotated gene; the remaining 2451 AS were classified as genic AS. From the latter group, 673 AS were classified as type 1 AS when both AS borders where flanked by at least one actively transcribed genes (distance of both AS borders to the closest transcribed (TPM>3) gene body was <20 kb), and 1026 AS were classified as type 2 AS when only one AS border was associated to a transcribed gene (see also Table 1). In order to assess the role of H4H20me3 mark on AS specification, we also classified as H4K20me3-containing non-genic AS, where the input normed H4K20me3 relative read frequency was above the genome mean value by more than 1.5 standard deviation (also estimated over the whole genome). This resulted in 154 non-genic AS with H4K20me3 higher than genome average and 242 non-genic AS with H4K20me3 signal lower than genome average.

### Comparison of ChIP relative read frequencies to transcription data

Gene containing bins were determined and overlapping genes removed from the analysis. For cumulative analysis, we only worked with genes larger 30kb, and assigned the gene expression levels in TPM accordingly. Genes were either aligned at their transcriptional start site (TSS) or their transcriptional termination site (TTS) and the corresponding ChIP read frequencies were calculated in a 30kb window around the site.

### Comparison of ChIP relative read frequencies to replication timing

For identification of RTDs in Raji cells, we used the early to late replication timing ratio determined by Repli-seq^43^. We directly worked from the precomputed early to late log-ratio from supplementary file GSE102522_Raji_log2_hg19.txt downloaded from GEO (accession number GSE102522). The timing of every non-overlapping

10 kb bin was calculated as the averaged log2 (early/late) ratio within the surrounding 100 kb window. Early RTDs were defined as regions where the average log-ratio > 1.6 and late RTDs as regions where the average log-ratio < -2.0. These RTDs were used to classify ChIP read relative frequencies calculated in 10 kb bins as early or late replication timing. Bins overlapping gene extended by 10 kb on both sides were removed from the analysis to avoid effects of gene activity on ChIP signals.

### Data access

Data has been deposited to the European Nucleotide Archive (ENA, https://www.ebi.ac.uk/ena). OK-seq data in Raji cells is available under the accession numbers PRJEB25180 (study) and SAMEA104651899 (sample accession, 5 replicates). Raji RNA-seq data is available under the accession number PRJEB31867 (study) and SAMEA5537240, SAMEA5537246, and SAMEA5537252 (sample accession per replicate). Raji ChIP-seq data was deposited under the accession number PRJEB32855.

## Acknowledgements

We thank Tobias Straub for initial help with bioinformatical analyses, Torsten Krude and Till Bartke for critical comments on the manuscript. A.S. was supported by the Deutsche Forschungsgemeinschaft (SFB 1064 TP05), SPP1230 and by the HELENA graduate school of the Helmholtz Zentrum München. B.A. and O.H were supported by the Agence Nationale de la Recherche (ANR-15-CE12-0011). O.H. was supported by the Ligue Nationale Contre le Cancer (Comité de Paris), the Association pour la Recherche sur le Cancer, the Fondation pour la Recherche Médicale (FRM DEI201512344404), the Cancéropôle Ile-de-France and the INCa (PL-BIO16-302), and the program “Investissements d’Avenir” launched by the French Government and implemented by the ANR (ANR-10-IDEX-0001-02 PSL*Research University). W.H. was supported by Deutsche Forschungsgemeinschaft (SFB1064/TP A13, SFB-TR36/TP A04), Deutsche Krebshilfe (grant number 70112875), and National Cancer Institute (grant number CA70723).

## Author contributions

N.K. designed and performed the majority of experiments; A.B. performed the RNA-seq experiment and TPM analysis; X.W. performed OK-seq experiments, S.K. and H.B. generated the sequencing library and sequencing, W.H. designed RNA-seq experiments; O.H. developed OK-seq, B.A supervised bioinformatic analyses; B.A. and N.K. performed bioinformatic analyses; A.S. proposed and designed the project and experimental systems; N.K. and A.S. wrote the manuscript with comments from O.H and B.A.; All the authors read and approved the manuscript.

## Competing Interests statement

The authors declare no competing interests.

## Supplementary tables

**Supplementary Table 1:** Proportion of genes significantly depleted from ORC/ Mcm2-7. A total of 1,941 genes met the criteria of transcriptional activity (TPM > 3), gene size larger 30 kb and no adjacent genes within 15 kb. We calculated the proportion of genes where the mean relative read frequency within the gene was significantly (p < 0.05) reduced compared to the upstream region (excluding TSS +/-3 kb).

| | | Total genes | $p < 0.05$ | % |
| --- | --- | --- | --- | --- |
| *pre* | Orc2 | 1,941 | 865 | 44.6 |
|  | Orc3 |  | 850 | 43.8 |
|  | Mcm3 |  | 1,460 | 75.2 |
|  | Mcm7 |  | 1,131 | 58.3 |
| *post* | Orc2 |  | 651 | 33.5 |
|  | Orc3 |  | 727 | 37.5 |
|  | Mcm3 |  | 554 | 28.5 |
|  | Mcm7 |  | 324 | 16.7 |

**Supplementary Table 2:** Characterization of H4K20me3 and H4K20me1 peaks determined by MACS2 broad peak calling.

|  | Number of peaks<br>(peaks > 1 kb) | Mean peak size<br>[kb] | Peak size range<br>[kb] |
| --- | --- | --- | --- |
| H4K20me3 | 16,852 (12,251) | 3.5 | 0.2-105.1 |
| H4K20me1 | 12,264 (6,277) | 5.5 | 0.2-182.5 |

**Supplementary Table 3:**
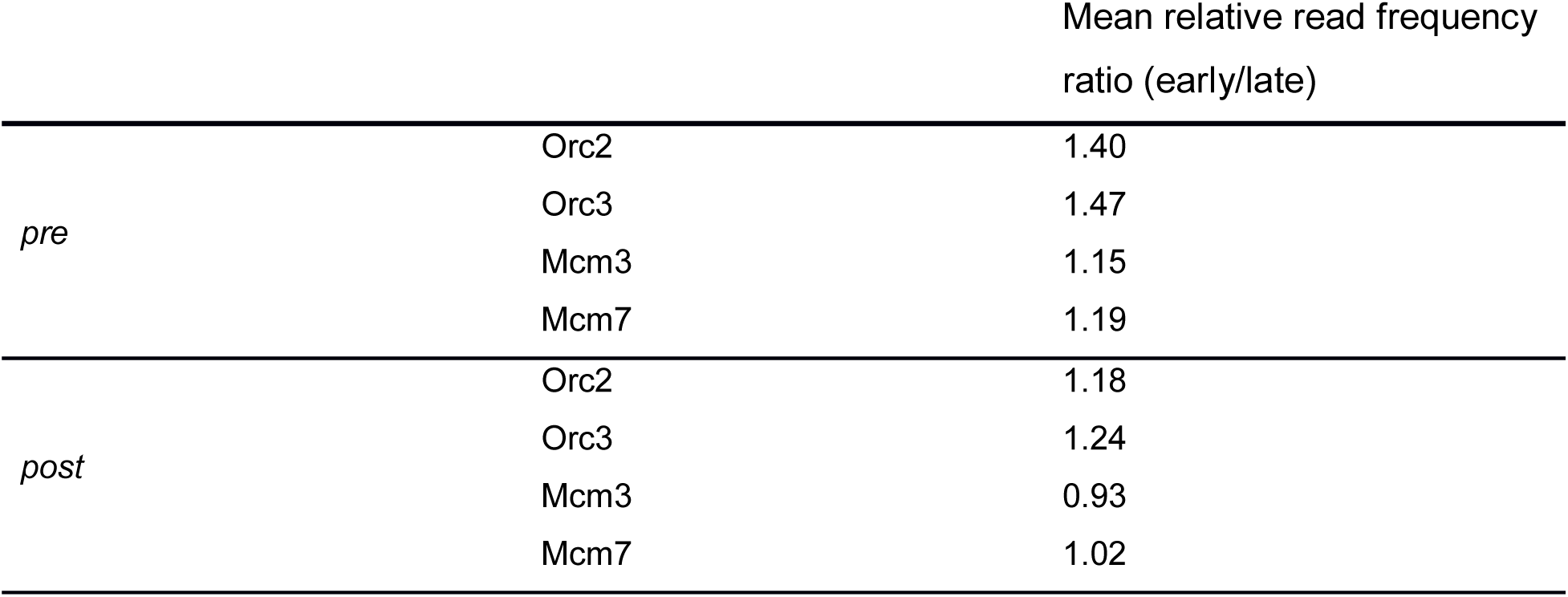
Ratio of ChIP mean relative read frequencies in early vs. late RTDs. Calculated in 10 kb bins. All annotated genic regions were removed ± 10 kb.

## Supplementary Figure Legends

**Supplementary Figure 1:**
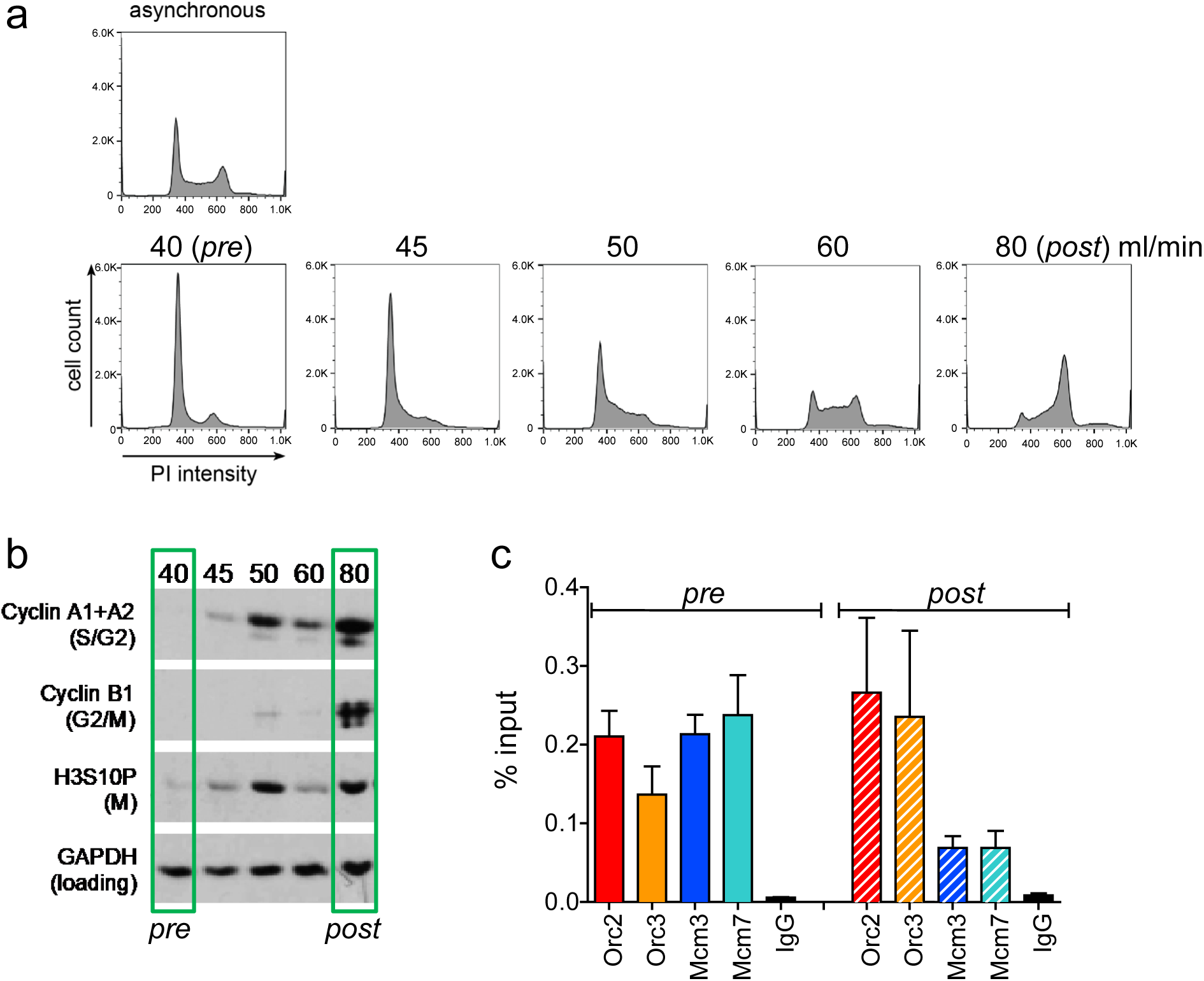
Experimental validation of cell cycle fractionation and ChIP quality. a) Example DNA content (Propidium Iodide) staining followed by FACS of logarithmically growing Raji (top) cells after cell cycle fractionation by centrifugal elutriation (increasing counter flow rates indicated above each profile). b) Western Blot analyses of the single fractions detecting Cyclin A (S/G2), Cyclin B (G2/M), H3S10P (M) and GAPDH. c) qPCR validation of Orc2, Orc3, Mcm3 and Mcm7 enrichment at the EBV latent origin *oriP* DS element. Representation in % input. Isotype IgG was used as control.

**Supplementary Figure 2:**
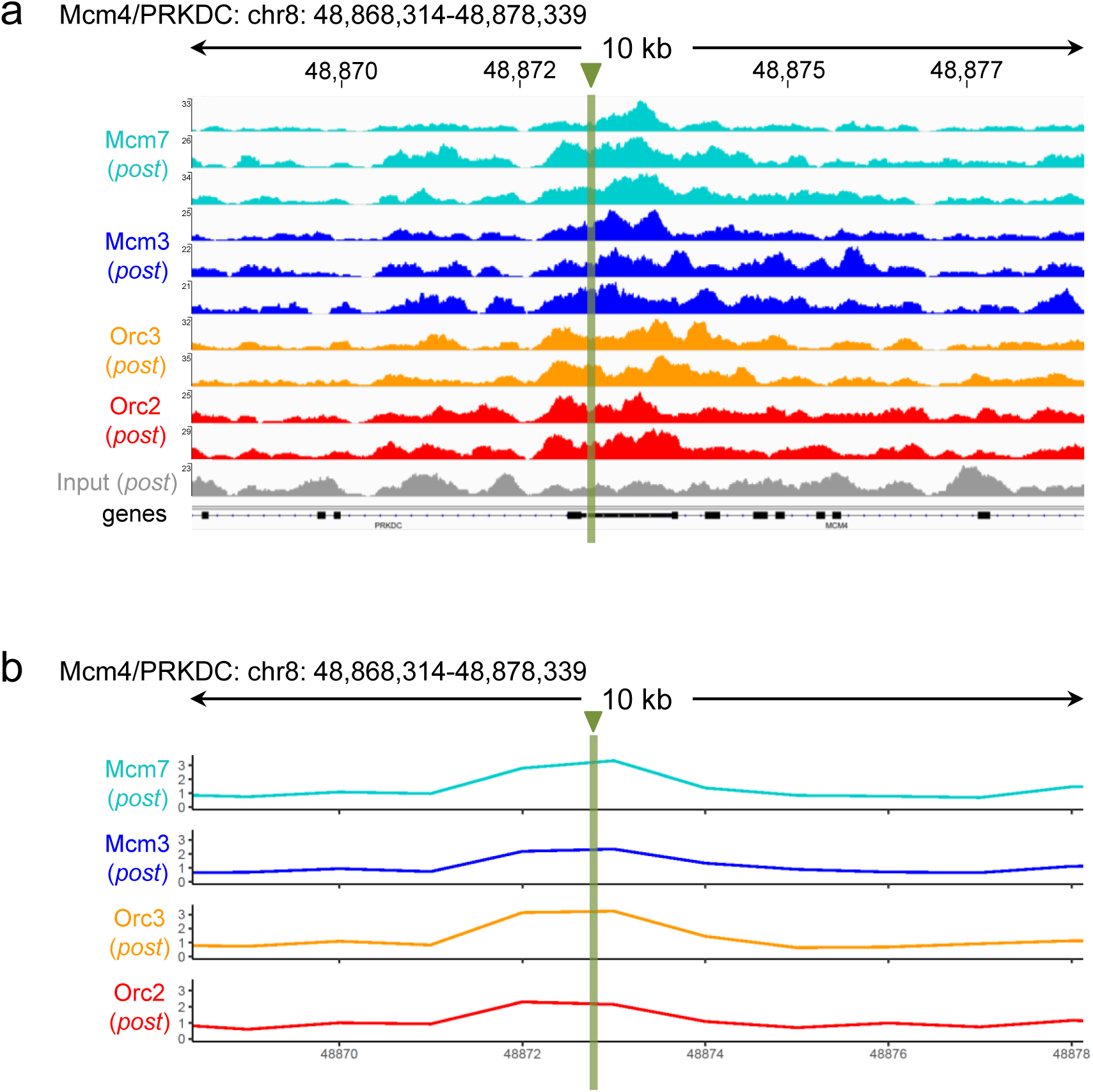
ORC/ MCM ChIP-seq profiles at the MCM4/ PRKDC origin before and after moderate averaging in *post*-fractions. a) Sequencing pileup IGB visualization of two samples of Orc2, Orc3 and three samples of Mcm3, Mcm7, plotted against the input at the Mcm4/PRKDC origin (*post*-fraction, chr8: 48,868,314 - 48,878,339).Track heights represent raw read depth. b) The ORC/ MCM ChIP-seq profile after 1 kb binning at the Mcm4/PRKDC origin (*post*-fraction). The reads of replicates were summed and normalized by the total ChIP read frequency followed by input division. Y-axis represents the resulting relative read frequency. The position of origin is indicated as green line.

**Supplementary Figure 3:**
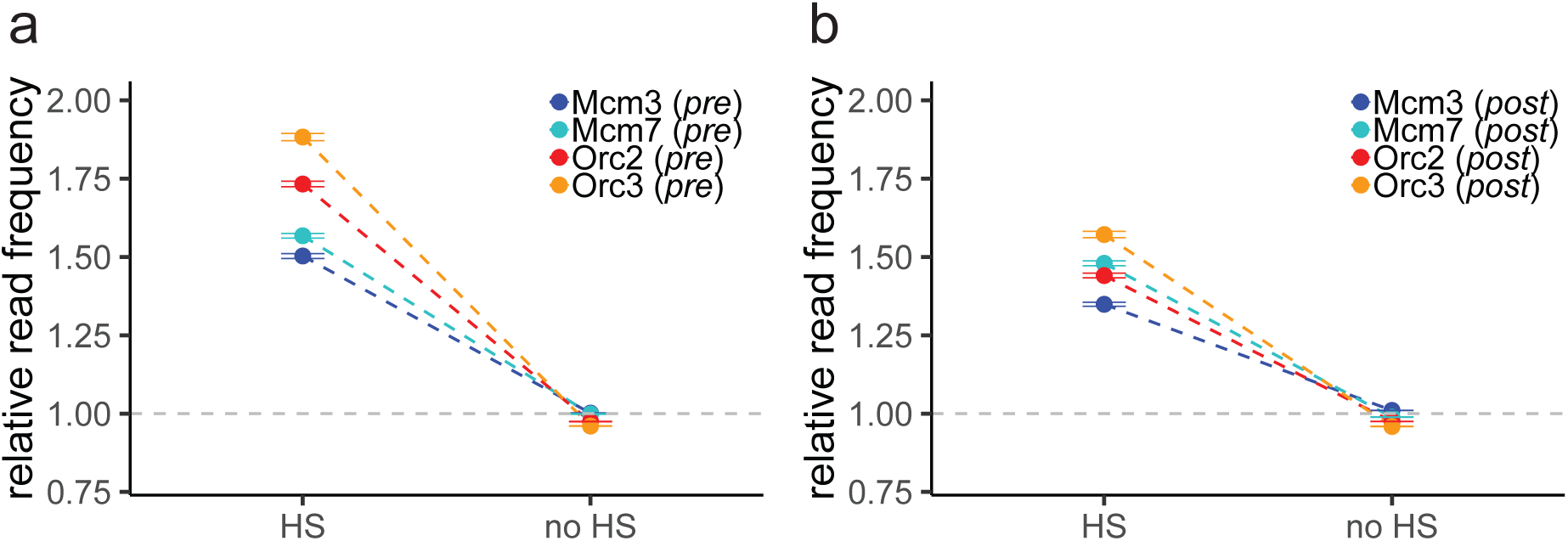
ORC/ MCM binding is confirmed at DNase HS sites. Mean ORC/ MCM relative read frequencies (± 2 x SEM) in relation to DNase hypersensitivity a) of the *pre*-fraction and b) of the *post*-fraction. Only HS sites larger 1 kb were considered. The dashed grey horizontal line indicates relative read frequency 1.0 for orientation.

**Supplementary Figure 4:**
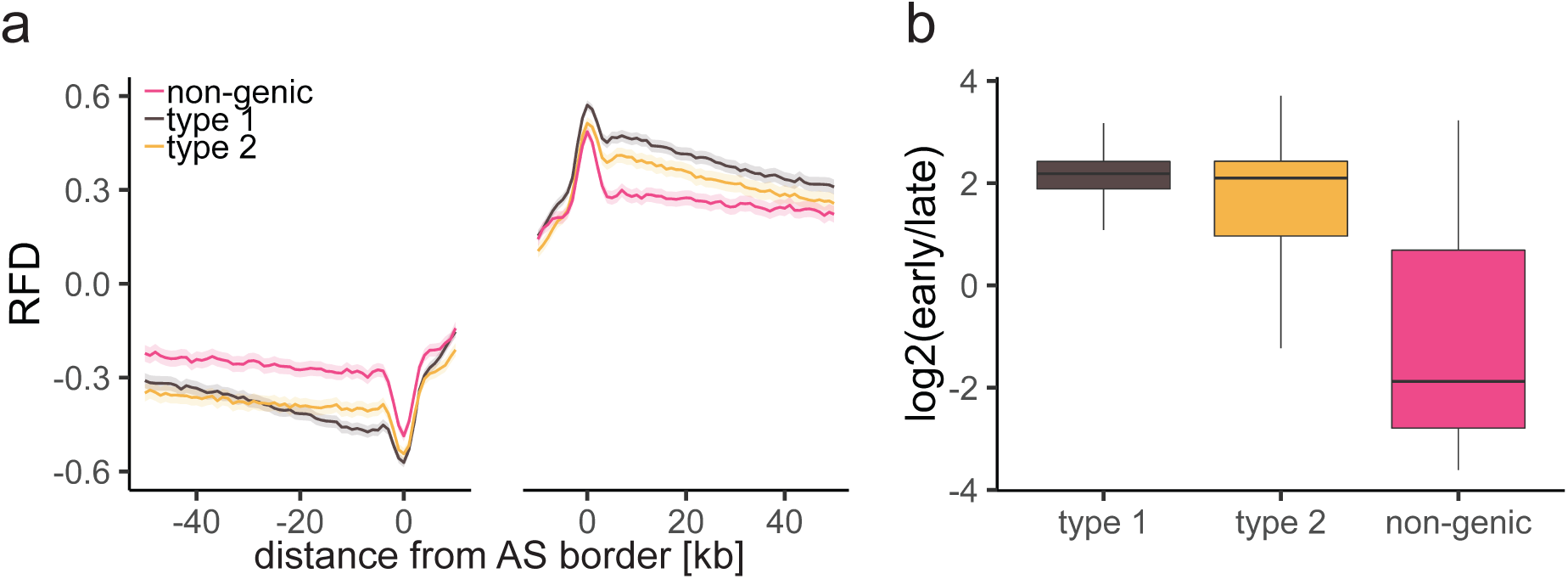
Characterization of different AS types. a) RFD of different AS types plotted at AS borders ± 2 x SEM (lighter shadows). b) Replication timing ratio log2 (early/late) was assigned to type 1, type 2, and non-genic AS and represented as boxplot.

**Supplementary Figure 5:**
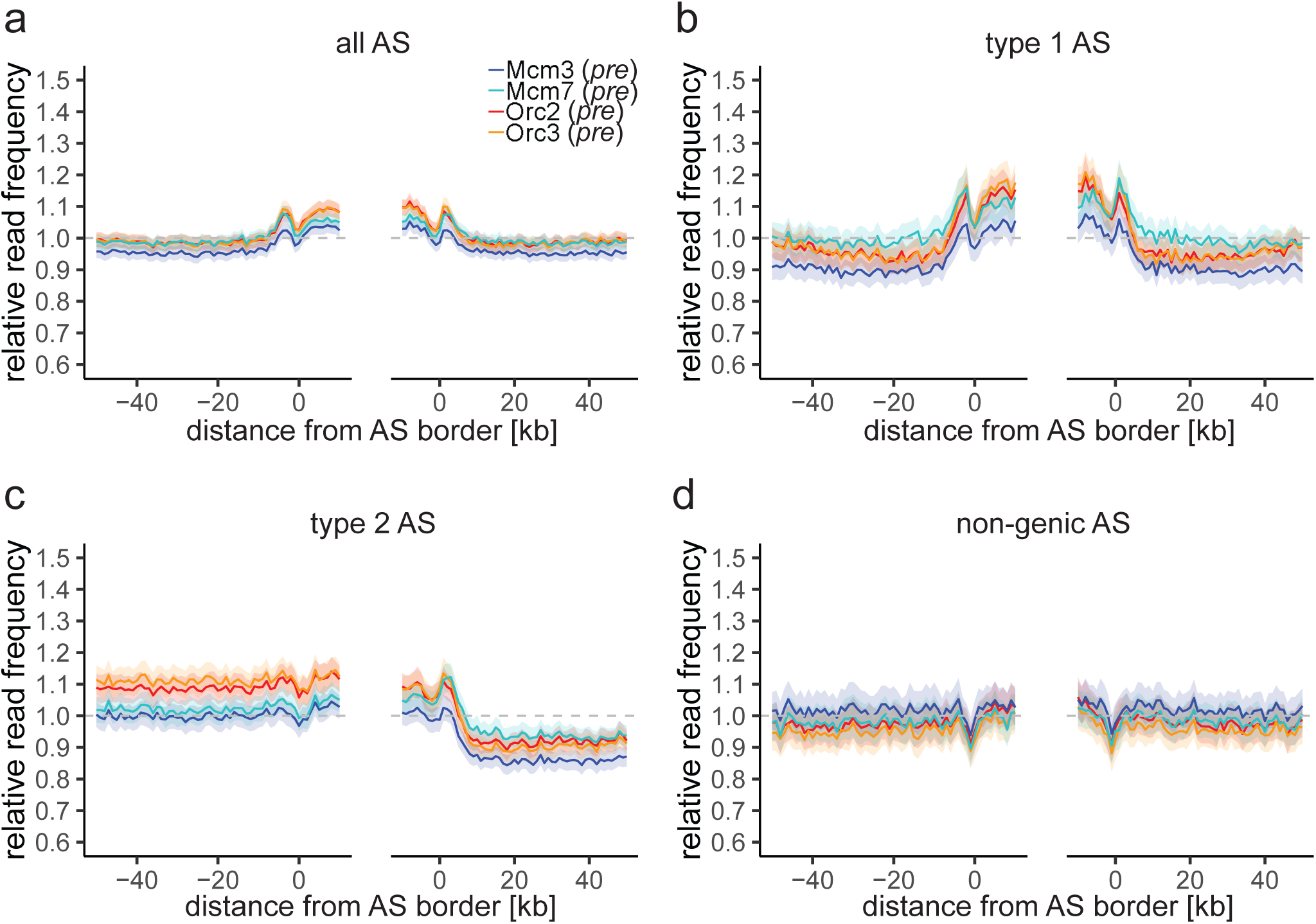
ORC/ MCM enrichment decreases within AS in the post-fraction. a)-d) Average relative ChIP read frequencies of Orc2, Orc3, Mcm3, and Mcm7 *post*-fractions at AS borders of a) all AS, b) type 1 AS with transcribed genes at both AS borders, c) type 2 AS with transcribed genes oriented at the right AS border, and d) non-genic AS in gene deprived regions. The mean of ORC and MCM frequencies are shown ± 2 x SEM (lighter shadows). The dashed grey horizontal line indicates relative read frequency 1.0 for orientation.

**Supplementary Figure 6:**
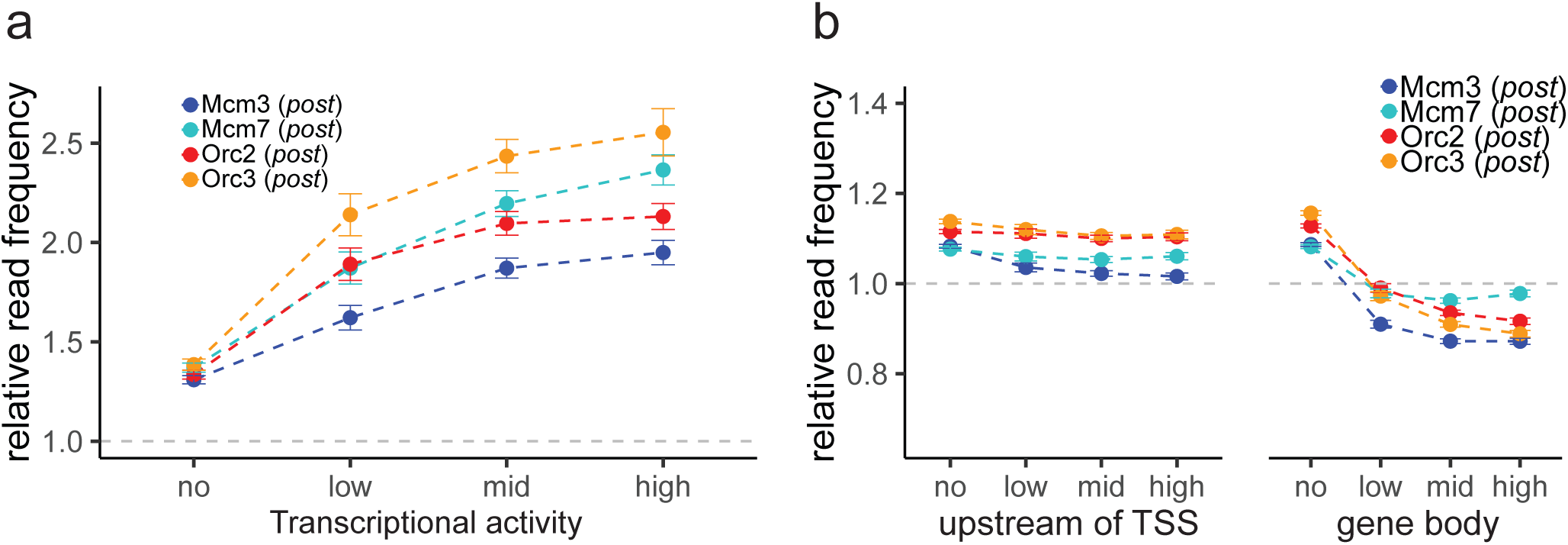
ORC/ MCM ChIP read frequencies at TSS and upstream and downstream of TSS in the *post*-fraction. a) ORC/ MCM (*post*) relative read frequencies at TSS dependent on transcriptional activity. Transcriptional activity was classified as: no (TPM < 3), low (TPM 3-10), mid (TPM 10-40), high (TPM > 40). b) ORC/ MCM (*post*) relative read frequencies upstream of TSS and in the gene body dependent on transcriptional activity (TSS ± 3 kb removed from analysis). Error bars correspond to ± 2 x SEM. The dashed grey horizontal line indicates relative read frequency 1.0 for orientation.

**Supplementary Figure 7:**
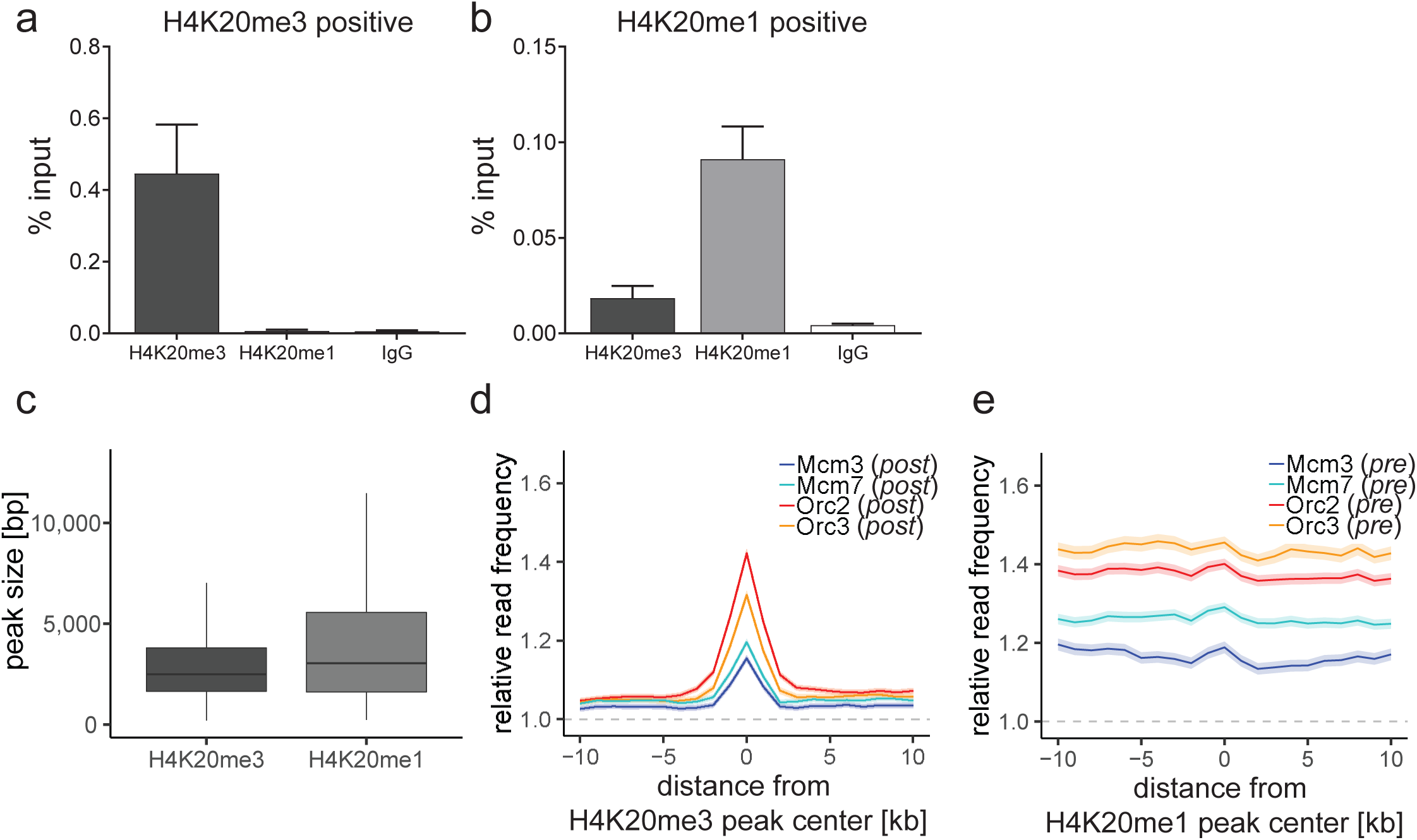
Characterization of H4K20 methylation profiles. a) and b) qPCR validation of H4K20me3 and H4K20me1 enrichment after ChIP at a) an H4K20me3 positive locus and b) an H4K20me1 positive locus. Representation in % input. Isotype IgG was used as control. c) Boxplot of H4K20me3 and H4K20me1 peak size (in bp) distribution. d) Average ORC/ MCM relative read frequencies (*post*-fraction) at H4K20me3 peaks (> 1 kb). e) Average ORC/ MCM relative read frequencies (*pre*-fraction) at H4K20me1 peaks (> 1 kb).

